# Automatic and unbiased segmentation and quantification of myofibers in skeletal muscle

**DOI:** 10.1101/2021.01.20.427432

**Authors:** Ariel Waisman, Alessandra Norris, Martín Elías Costa, Daniel Kopinke

## Abstract

Skeletal muscle has the remarkable ability to regenerate. However, with age and disease muscle strength and function decline. Myofiber size, which is affected by injury and disease, is a critical measurement to assess muscle health. Here, we test and apply Cellpose, a recently developed deep learning algorithm, to automatically segment myofibers within murine skeletal muscle. We first show that tissue fixation is necessary to preserve cellular structures such as primary cilia, small cellular antennae, and adipocyte lipid droplets. However, fixation generates heterogeneous myofiber labeling, which impedes intensity-based segmentation. We demonstrate that Cellpose efficiently delineates thousands of individual myofibers outlined by a variety of markers, even within fixed tissue with highly uneven myofiber staining. We created a novel ImageJ plugin (LabelsToRois) that allows processing multiple Cellpose segmentation images in batch. The plugin also contains a semi-automatic erosion function to correct for the area bias introduced by the different stainings, identifying myofibers as accurately as human experts. We successfully applied our segmentation pipeline to uncover myofiber size differences between two different muscle injury models, cardiotoxin and glycerol. Thus, Cellpose combined with LabelsToRois allows for fast, unbiased, and reproducible myofiber quantification for a variety of staining and fixation conditions.

## Introduction

In many tissues, wound healing and regeneration depends on stem cells to replace the lost or damaged cells. For example, skeletal muscle has the remarkable ability to fully recover from injury due to a dedicated muscle stem cell population (MuSCs)^1–4^. Injury activates the MuSCs to replace damaged or lost myofibers. Skeletal muscle contains a second type of stem cell, called fibro/adipogenic progenitors (FAPs). FAPs are mesenchymal stem cells that work with MuSCs to regenerate skeletal muscle^5–12^. Following an acute injury, FAPs transiently expand and promote MuSC differentiation. In chronic diseases, however, muscle regeneration fails and FAPs produce scar tissue and differentiate into adipocytes^5,6,10,11,13–16^. This fatty fibrosis of muscle is a prominent feature of chronic muscle diseases such as Duchenne muscular dystrophy (DMD), sarcopenia, the agerelated loss of skeletal muscle and strength, obesity, and dia betes^13, 17, 18, 18–26^.

A critical measurement to evaluate muscle health or the ability to recover from injury is the cross-sectional area (CSA) of myofibers^8 27–30^,. Myofiber CSA can increase due to hypertrophy, where myofibers become larger, for example, after consistent exercise or during the initial phase of muscular dystrophies like DMD^31–33^. Reduced myofiber CSA indicates impaired regeneration after an acute injury and contributes to the progressive decline in muscle strength seen with age and disease^34^. Unbiased and rigorous quantification of myofiber CSA provides, therefore, crucial information when studying the cellular and molecular mechanisms of muscle regeneration as well as assessing potential therapeutics to prevent the decline in muscle mass and strength with age and disease.

Quantification of myofiber CSA is routinely done by snap-freezing muscle tissues followed by immunohistochemistry for markers that outline individual myofibers such as Laminin. This method provides robust and homogeneous staining of myofiber’s contours and can be easily used to accurately delineate hundreds of myofibers using the thresholding function in ImageJ. In fact, several sophisticated ImageJ or Matlab-based plugins have been developed over the years to allow for semi-automatic quantification of myofibers^35–39^. However, snap-freezing is not compatible with immunostainings for certain epitopes that require prior fixation with fixatives such as paraformaldehyde (PFA). For instance, in our experience, prior fixation is crucial to preserve the morphology of individual adipocytes, especially the lipid droplets, that infiltrate muscle. As the functional relevance of intramuscular fat is still unclear, it is important to preserve adipocytes within the muscle to study their function. Additionally, we noticed that some cellular structures and organelles, such as filopodia and primary cilia, require fixation to preserve their integrity. Cilia are small organelles that extrude from the cell and act as a cellular antenna that sense extracellular cues^6^. We have previously shown that cilia control the differentiation of FAPs into adipocytes in skeletal muscle^6^ and white adipose tissue^40^ by controlling both anti- and pro-adipogenic cues, respecitvly. In addition, FAP cilia are critical for controlling muscle regeneration after an acute injury and maintaining myofiber size in a mouse model of DMD, the mdx mouse^6^. However, without prior fixation cilia are undetectable. To circumvent this challenge, the muscle group of choice can be either cut in half and divided into two samples or the experiment has to be repeated with a new set of animals resulting in reduced efficiency, increased animal numbers, and a missed opportunity to study different cellular processes at the same Z plane. The major disadvantage of prior fixation is that PFA causes uneven staining and a reduced signal-to-noise ratio of most commonly used myofiber markers. This presents a major challenge for using PFA-fixed muscle sections to quantify large numbers of myofibers and especially prevents the use of the previously developed CSA plugin tools.

Here, we tested and applied Cellpose to automatically segment individual myofibers using a variety of staining and fixing conditions. Cellpose is a deep-learning segmentation algorithm that was recently developed as an efficient way to segment a variety of cells stained for different markers^41^. Cellpose allowed us to readily and efficiently identify thousands of myofibers regardless of dye or antibody used. Most importantly, this tool was able to robustly segment myofibers even with suboptimal myofiber labeling caused by PFA-fixation. However, depending on the staining used, the outline of the myofibers was larger than expected, significantly affecting CSA or other area-related measurements. To circumvent this, we developed a novel ImageJ plugin, LabelsToROIs, which allows processing Cellpose generated label images within ImageJ. It overlays the Cellpose-generated segmentations with the raw file followed by automatic quantification of the crosssectional area (CSA) of each myofiber or other desired measurements. We also integrated a segmentation erosion tool to counter the above-mentioned size bias, which can be as high as 30%. We successfully validated the use of Cellpose together with our ImageJ plugin in different muscle regeneration and disease paradigms. Thus, the use of Cellpose in combination with our ImageJ plugin saves time, reduces bias, and allows for quantification of poorly stained images. Furthermore, it is applicable to a wide variety of staining conditions, and, with the detailed step-by-step instructions provided, can be widely adopted by users without programming skills.

## Results

### Cellpose efficiently identifies myofibers regardless of staining or fixation condition

The preservation of many cellular structures requires immediate fixation of tissue biopsies. For example, we noticed that filopodia, primary cilia and adipocytes in skeletal muscle tissue are highly sensitive to fixation. We previously discovered that FAPs are the main ciliated cell type in muscle^6^. Confirming our previous findings, primary cilia, marked by the small GTPase ARL13B, are readily detected on PDGFRA expressing FAPs in paraformaldehyde (PFA) fixed tibialis anterior (TA) muscle (Fig. 1a). However, without fixation, we failed to detect ARL13B^+^ cilia (Fig. 1a). We were also unable to detect cilia using acetylated tubulin, a marker for the ciliary axoneme (data not shown) in unfixed tissue sections. In addition, PDGFRA expression was less robust in snap-frozen tissues preventing the preservation of the intricate cellular processes and filopodia of FAPs (Fig. 1a). Glycerol (GLY) injection into the TA muscle causes massive fat infiltration similar to what is seen in chronic neuromuscular disease and sarcopenia^5,6,11,28,42–44^. The resulting adipocytes can be robustly stained for markers such as PERILIPIN 21 days post-GLY injury (Fig. 1b). However, adipocyte morphology is severely compromised in unfixed tissue hindering faithful quantification of absolute adipocyte numbers (Fig. 1b). Successful muscle regeneration post-injury is evaluated by determining the CSA of the regenerated myofibers. For this, muscle sections are routinely stained for LAMININ, a basal lamina protein that outlines individual myofibers (Fig. 1c). However, LAMININ staining is highly sensitive to fixation (Fig. 1c). Thus, while standard thresholding in ImageJ allowed for simple fiber segmentation of snap-frozen muscle sections, fixation resulted in significant background preventing the use of fixed muscle tissues for evaluating crosssectional area measurements (Fig. 1c).

**Figure 1.**
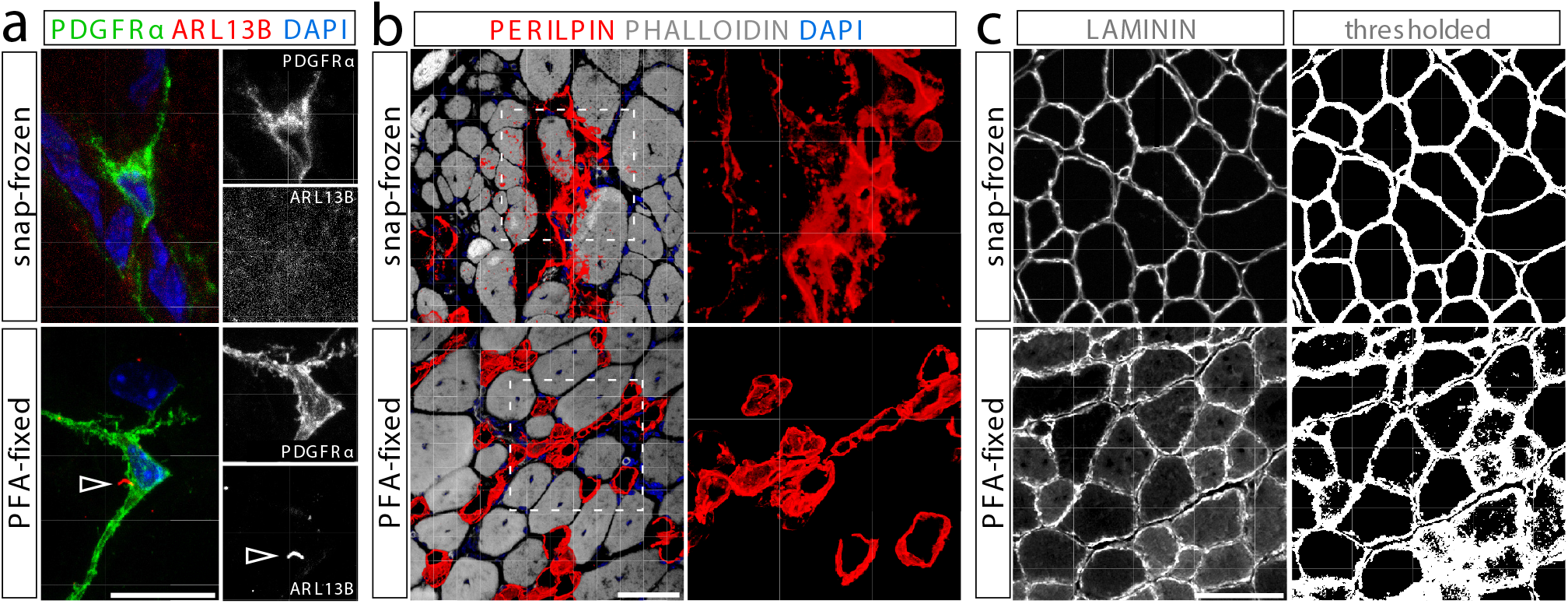
Fixation of muscle tissues is required to preserve certain cellular structures. (a) Cilia, marked by ARL13B (red), can only be detected on FAPs, labeled by PDGFR (green), in PFA-fixed (bottom) but not snap-frozen (top) tibialis anterior muscle sections. Scale bar is 10 µm. (b) The morphology of PERILIPIN-expressing adipocytes (red) is severely compromised without prior fixation. Scale bar is 50 µm. (c) LAMININ (gray) perfectly outlines myofibers in snap-frozen tissue (top) enabling simple myofiber segmentation using ImageJ thresholding. However, fixation impacts LAMININ immunoreactivity preventing myofiber segmentation. Scale bar is 250 µm.

A novel deep-learning algorithm, called Cellpose^41^, was recently designed to robustly and automatically segment individual cells from dense cell clusters. The output of Cellpose are labeled images, an image data format where the segmentations are stored and can be eventually converted to the ImageJ regions of interest (ROIs) for subsequent analyses (see below for more details). We evaluated the performance of the Cellpose algorithm in analyzing cross-sections of the TA muscle stained for different markers (Fig. 2a). We first determined whether this method could efficiently identify myofibers in cross-sections of snap-frozen TA muscles stained for LAMININ, the gold standard in the muscle field. Snap freezing muscle tissue allows for high-quality LAMININ images, and, thus, it is more amenable for automatic segmentation. As seen in Fig. 2b, Cellpose allowed for the identification of thousands of myofibers within all size ranges with great accuracy, without any necessary pre-processing of the images. After these promising results, we next assessed if this algorithm could also identify myofibers in PFA-fixed tissue. As described above, fixation causes a more heterogeneous LAMININ staining, with areas of different LAMININ intensities and different internal unspecific background within the fibers. Due to this complexity and heterogeneity, many of the currently available automatic segmentation methods that rely on fluorescent intensity thresholds fail to segment PFA-fixed muscle. Remarkably, Cellpose could efficiently identify the myofibers in these complex images with results highly similar to the gold standard (Fig. 2c). Similar results were obtained for the gastrocnemius muscle (Fig. S1).

**Figure 2.**
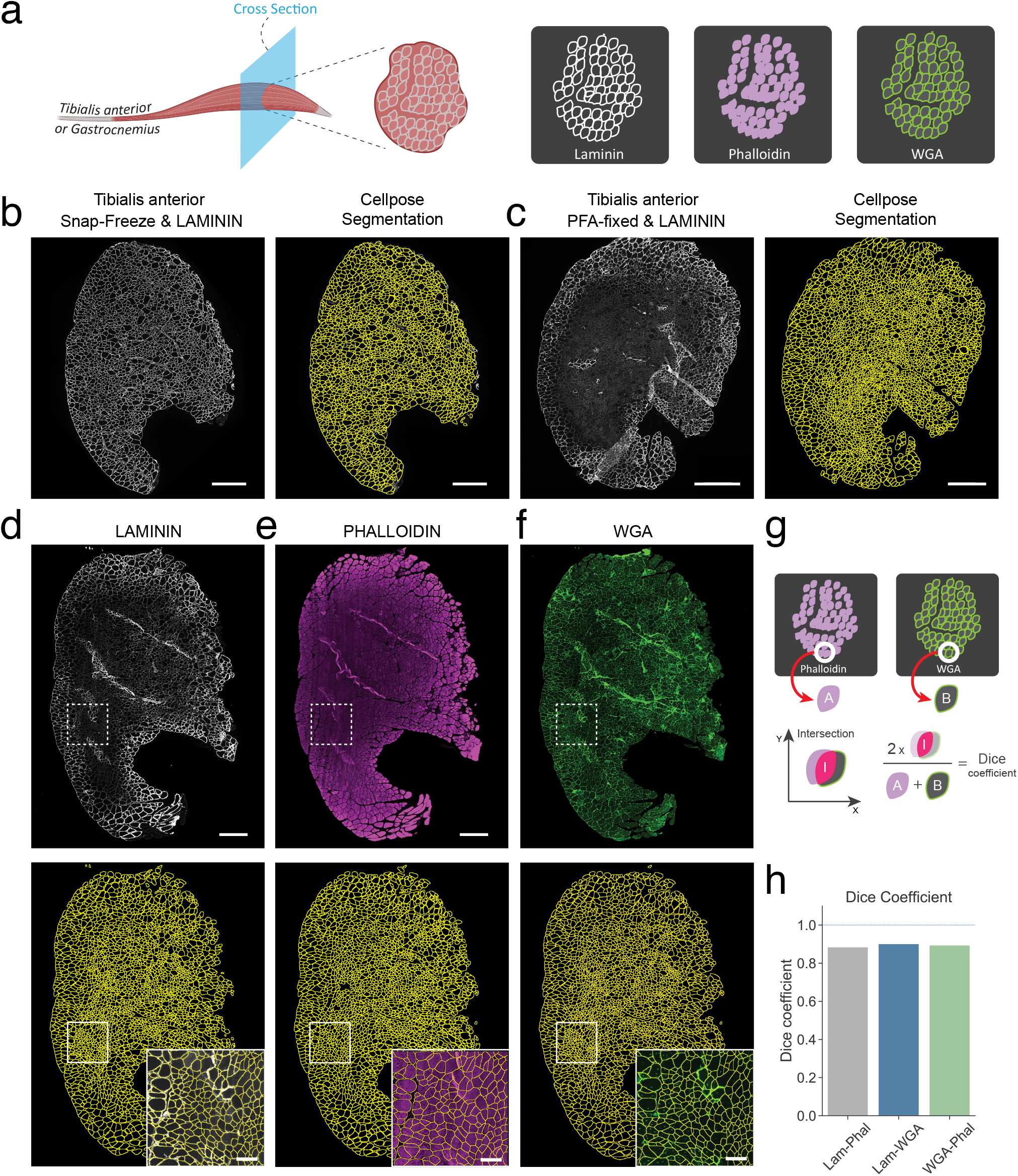
Myofiber segmentation using Cellpose. (a) Schematic of the experimental design. (b) Cross section of a snapfrozen tibialis anterior muscle stained for LAMININ. Right, original image. Left, Cellpose segmentation results. Scale bar, 500 µm. (c) Cross section of a PFA-fixed tibialis anterior muscle stained for LAMININ. Right, original image. Left, Cellpose segmentation results. Scale bar, 500 µm. (d) Cross section of a PFA-fixed tibialis anterior muscle stained for LAMININ and with Phalloidin and WGA. Top, original images. Bottom, result of Cellpose segmentation. Inset, detail of the specified region. Scale bar within inset, 500 µm. (g) Diagram depicting the calculation of the Dice coefficient. For each segmented object in the two label images, the Dice coefficient is the result of two times the area of the intersection divided by the sum of the areas of both objects. (h) Mean Dice coefficient for all myofibers in the comparison between the indicated stains.

We next evaluated if Cellpose could identify myofibers in cross sections of muscle stained for other commonly used markers such as Phalloidin and wheat-germ agglutinin (WGA), which label the actin cytoskeleton and the cell membranes, respectively. After triple staining for LAMININ, Phalloidin and WGA dyes of PFA-fixed tibialis anterior cross sections, Cellpose could accurately identify thousands of myofibers in all three stains (Fig. 2d-f). Indeed, for this image, Cellpose identified approximately 3200 fibers for LAMININ, with Phalloidin and WGA closely matching this number, a task that would require days of repetitive and laborious work with manual segmentation. To further ascertain whether the segmentation for each of the fibers between the different stains was similar, we used the Dice coefficient^45–47^. As depicted in Fig. 2g, this metric provides a numerical quantification of the resemblance of two individual segmentations based on their overlap. A value of 1 corresponds to the same shape, size, and location, while 0 indicates no intersection between the segmentations. As can be seen in Fig. 2h the mean Dice coefficient for all the fibers between the different staining’s was approximately 0.9, indicating that the Cellpose algorithm generated highly similar segmentations for LAMININ, Phalloidin, and WGA. Thus, our results show that Cellpose can achieve a very precise and comprehensive identification of the myofibers, independent of the staining or fixation condition.

### Benchmarking of Cellpose across several computing platforms

We next assessed the performance of the Cellpose segmentation algorithm under different computer settings. Since the segmentation of each skeletal muscle image, although automatic, can take a considerable amount of computing time, we evaluated the effect of image downscaling on its overall segmentation performance. For a single-channel image, the only input that Cellpose takes besides the image itself is an approximate mean cell diameter in pixels, which the user must determine previously, for instance, by sampling a few representative myofibers within the image. We started our analysis with a PFA-fixed LAMININ cross-sectional image of TA muscle, with a resolution of 7059x11763 pixels (3199x5332 µm^2^) and an approximate myofiber diameter of 120 pixels (∼40 µm). We then created 4 downscaled versions of up to 20 times smaller, covering the range of 0.5x, 0.2x, 0.1x, and 0.05x, and then used Cellpose to generate the label images with the segmentations using different diameters (Fig. 3a,b).

**Figure 3.**
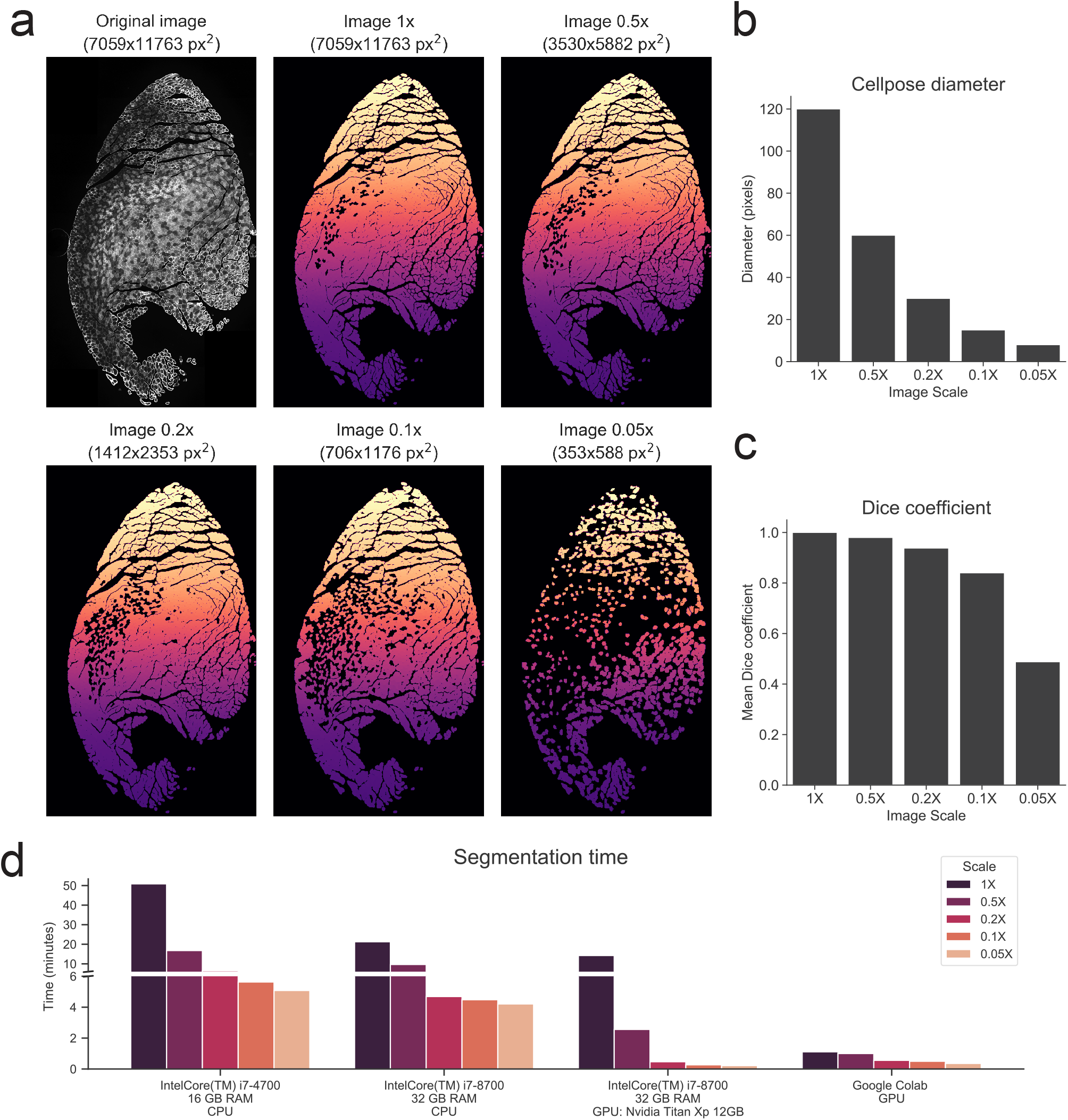
Benchmarking of Cellpose algorithm across different image sizes and hardware specifications. (a) Top-left, original full-sized PFA-fixed TA cross-section stained for LAMININ. The remaining images correspond to the Cellposegenerated label images when full-sized or downscaled versions of the original image where fed to the algorithm. (b) Myofiber diameter fed to the Cellpose algorithm for the full-sized or the downscaled images. Approximate myofiber diameter for the original image was manually determined. (c) Mean Dice coefficient for the comparison of the original 1X label image with itself or with the downscaled label images (see Materials and Methods). (d) Comparison of Cellpose segmentation time for the different image sized across different hardware platforms.

To objectively quantify Cellpose’s performance on the downscaled images, we calculated the mean Dice coefficient between the downscaled label images and the full-size 1X label image (Fig. 3c). These results show a gradual decrease in the segmentation quality with the amount of scaling. Importantly, when downscaled up to 5 times, reaching an approximate myofiber diameter of 30 pixels, the mean Dice coefficient compared to the full-size image was 0.94, thus showing a very acceptable segmentation quality. For the image downscaled 20 times, where the approximate myofiber diameter was 8 pixels, the mean Dice coefficient was 0.48, indicating an important decrease of the segmentation quality, as can be observed in (Fig. 3a).

We next evaluated Cellpose’s segmentation computing time for the differently scaled images. We did this using four different hardware configurations. For computers bearing a compatible Nvidia video card, Cellpose can be configured to run on the GPU, which can substantially decrease the computing time. We thus evaluated its performance on an Intel i7-8700 CPU with 32 GB of RAM bearing an Nvidia Titan Xp, running the different segmentations in the GPU or in the CPU-modes. We also evaluated the performance of a 2015 laptop computer with an i7-4700 CPU and 16 GB of RAM, running the segmentations using the CPU-mode only. Finally, we also made use of the Google colab online programming platform, which can be configured to run in a GPU mode. As can be seen in Fig. 3d, the resolution of the images greatly affected Cellpose’s computing time, with a clear advantage when running the segmentations on the GPU (for a detailed video tutorial, follow the link to the Github page in the Materials and Methods section). In the least powerful hardware, the full-size image took more than 50 minutes to segment. Therefore, we especially recommend the use of Google Colab, which enables rapid segmentation at full resolution without the need of a dedicated local computer. However, considering that the quality of downscaled images up to 30 pixels of cell diameter is reasonably acceptable, users should evaluate the compromise between the segmentation quality and computing time.

### Correction of segmentation-induced size bias

We noticed that Cellpose segmented the myofibers in such a way that it often included the LAMININ staining itself causing the myofiber area to appear artificially larger (Fig. 4a,b). To circumvent this problem, we eroded the myofiber’s segmentations pixel by pixel until LAMININ was largely excluded (Fig. 4c). We then quantified the mean CSA of LAMININ stained TA muscle cross-sections of 6 different mice with or without label erosion (Fig. 4d). These results show that a slight amount of label erosion can result in a significant difference in mean CSA, in this case, as much as 30%, reinforcing the importance of manual supervision.

**Figure 4.**
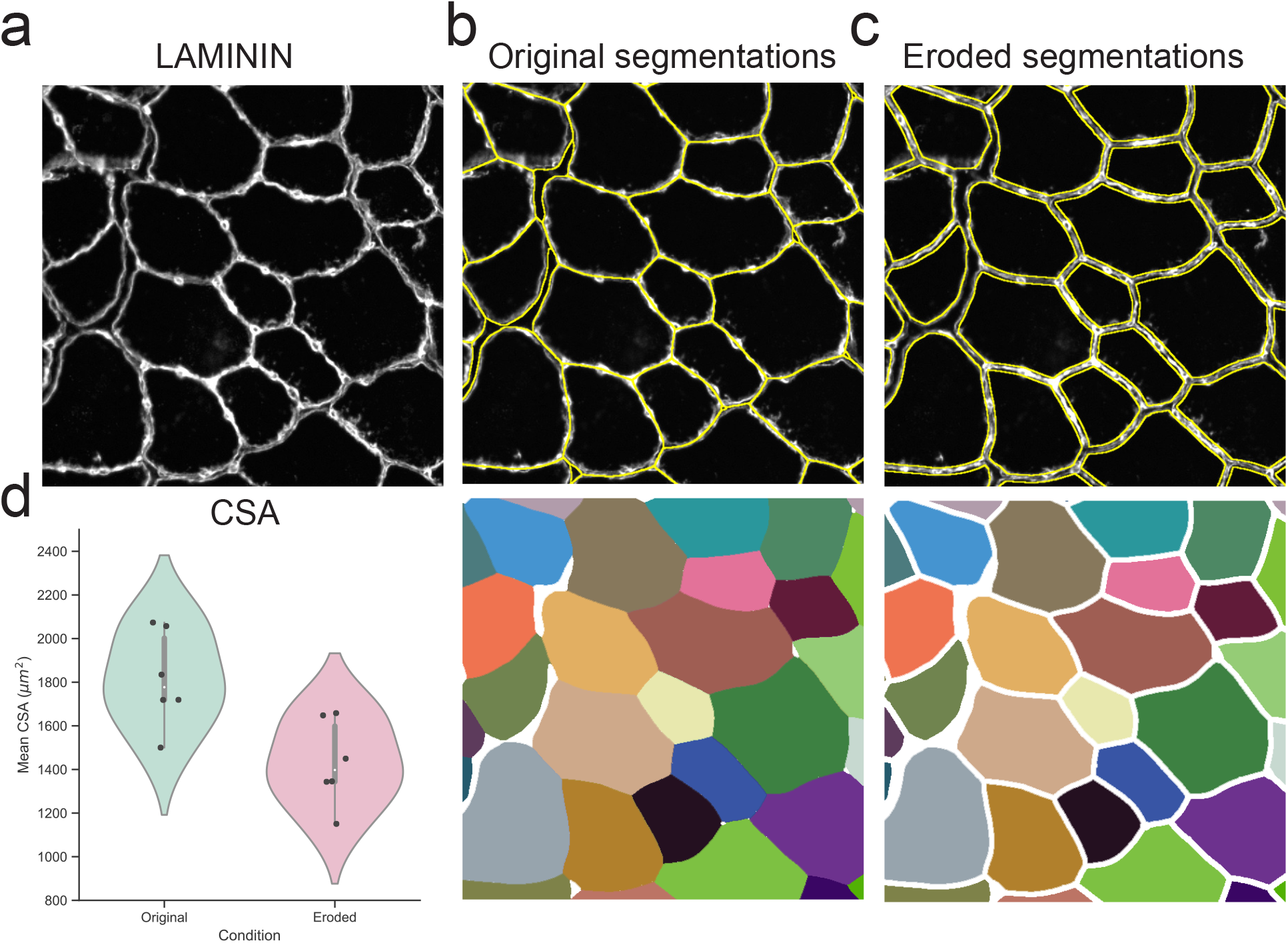
Erosion of Cellpose segmentations is needed for accurate CSA measurements. (a) Example of a snap-frozen LAMININ stained TA cross-section. (b) Cellpose segmentation of the image in (a) showing the overlaid segmentations in yellow (top) or the Cellpose-generated label image (bottom). (c) Segmentation results after the Cellpose label image was erorded by 4 pixels. (d) Mean CSA for all myofiber of LAMININ stained TA muscle cross-sectios of 6 different mice with or without a 4-pixel label erosion.

To further validate the use of Cellpose, we next analyzed how similar the segmentations generated by this algorithm (with or without erosion) were compared to manual segmentations generated by three human experts. To do this, PFA-fixed TA cross-sections were stained for LAMININ, Phalloidin, or WGA, and four images with approximately 50 myofibers each were obtained for each staining. Cellpose was then used to identify the myofibers in each image (CP) and, in parallel, these segmentations were eroded with a fixed number of pixels according to visual inspection (eroded CP, eCP). Additionally, three different human experts manually segmented the same images (M). Fig. 5a displays representative segmentations for each staining and condition.

**Figure 5.**
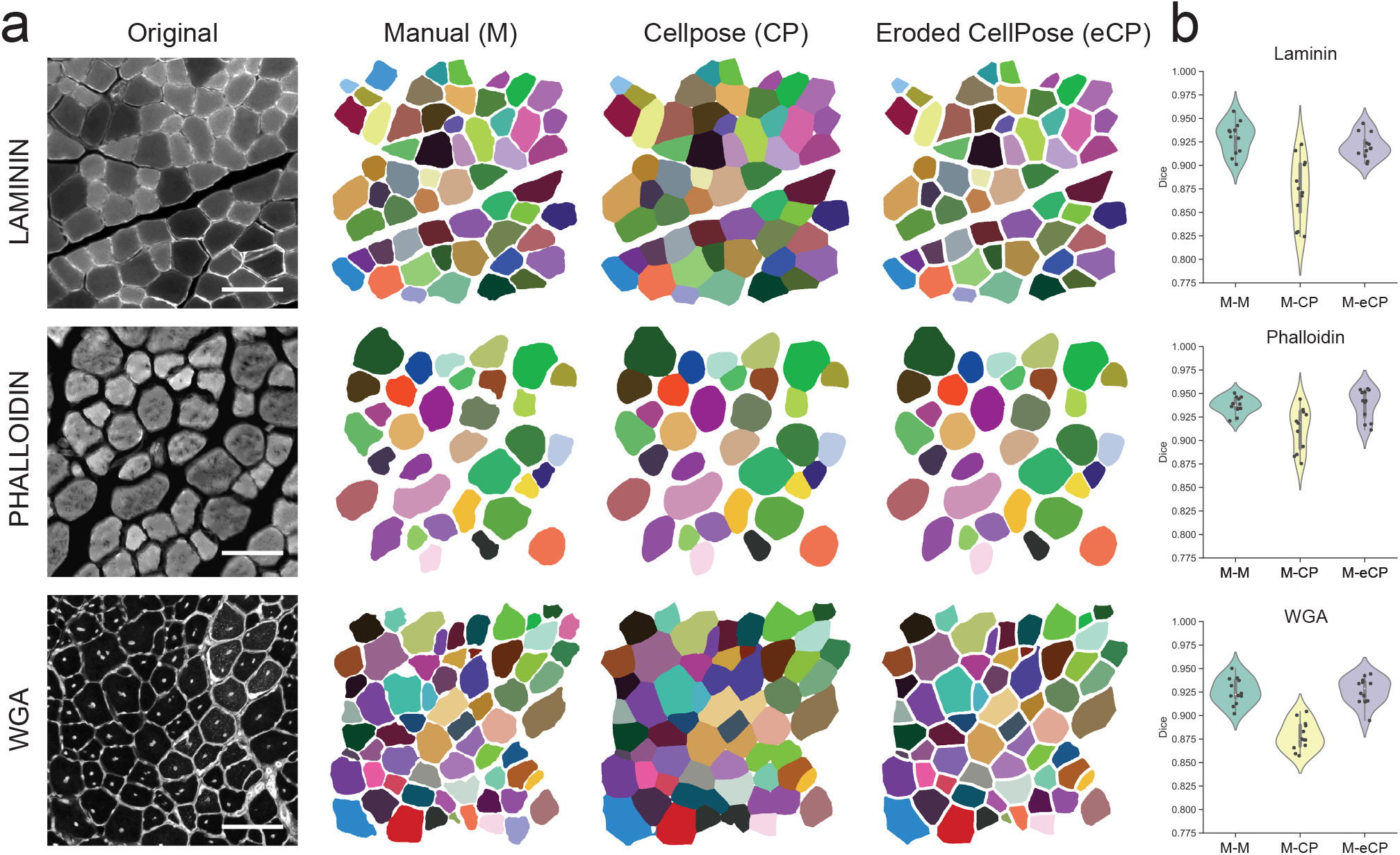
Eroded Cellpose segmentations are as accurate as human experts in delineating myofibers. (a) Left, example of cross-sectional TA images marked by LAMININ, Phalloidin or WGA and, right, their corresponding labeled images manually generated by a human expert (M), and the original (CP) or the eroded Cellpose segmentation (eCP). (b) A set of four images was analyzed for each of the three indicated stains. Three human experts manually segmented each of the images, while those same images were also segmented by Cellpose. Cellpose labels were additionally eroded by a fixed number of pixels according to visual inspection. The Dice coefficient was used to compare the different label images corresponding to the same original image. The violinplot shows the mean Dice coefficient for each human pair for a specific image (M-M), the comparison of human manual segmentations to their associated Cellpose segmentations (M-CP), or for human manual segmentations to their associated Cellpose eroded segmentations (M-eCP).

We used the Dice coefficient to objectively analyze the accuracy of the segmentations. Since different human experts can generate slightly different segmentations, we first calculated the distribution of Dice coefficients for each pair of human experts for the 4 images of each stain (M-M). This analysis showed, as expected, high values of Dice coefficient that ranged between 0.9 and 0.95, indicating an approximate 5 to 10% error rate in the manual segmentations (Fig. 5b). Next, we compared the segmentation of each expert to the one generated by Cellpose (M-CP), which shows that Cellpose’s segmentations, despite having high Dice coefficients, were less accurate than the manual segmentations. It is important to note that Cellpose’s LAMININ segmentation provided less accurate results than Phalloidin or WGA. Remarkably, after eroding the segmentations with a fixed number of pixels, the dice coefficient of the eroded labels compared to humans was practically indistinguishable from in-between humans (M-eCP). These results show that when combined with label erosion, Cellpose can identify myofibers as accurately as human experts.

### Development of the plugin LabelToRois to analyze and modify Cellpose segmentations in ImageJ

The current release of the Cellpose segmentation algorithm runs on Python and can be also used with a convenient user interface. As we previously mentioned, Cellpose’s segmentation results are in the format of labeled images, in which each segmentation has a specific pixel value that is used as an unequivocal identifier (see the accompanying tutorial in the material and methods section for more information about labeled images). This type of data storage, although quite common in the field of computer vision, is not widely used by users without programming skills. ImageJ is a powerful and user-friendly image analysis software widely adopted in the biological community. Within ImageJ, the Regions of Interest (ROIs) are an effective way to identify objects before making different measurements. However, there is currently no easy nor efficient way of transforming the information stored in label images to ImageJ ROIs for non-programmers while also allowing simultaneous ROI erosion.

To circumvent these issues and extend the power of Cellpose to as many users as possible, we developed an ImageJ plugin, LabelsToROIs, that takes label images as input and converts them into ROIs. Our plugin allows users to automatically erode the segmentations by a user-defined number of pixels while being able to visually inspect the resulting ROIs, delete, add, or modify them (Fig. 6). It includes functions to save the ROIs for future inspection, as well as to select which measurements to take (i.e. area, Ferret’s diameter, etc) on the original images and saving the results in convenient CSV tables. Furthermore, it also allows for batch analysis of multiple label images with minor intervention. Thus, this simple plugin allows users without programming skills to widely adopt this great segmentation tool. Importantly, this plugin works well for any kind of label image and, thus, its use is not limited to Cellpose nor the field of muscle.

**Figure 6.**
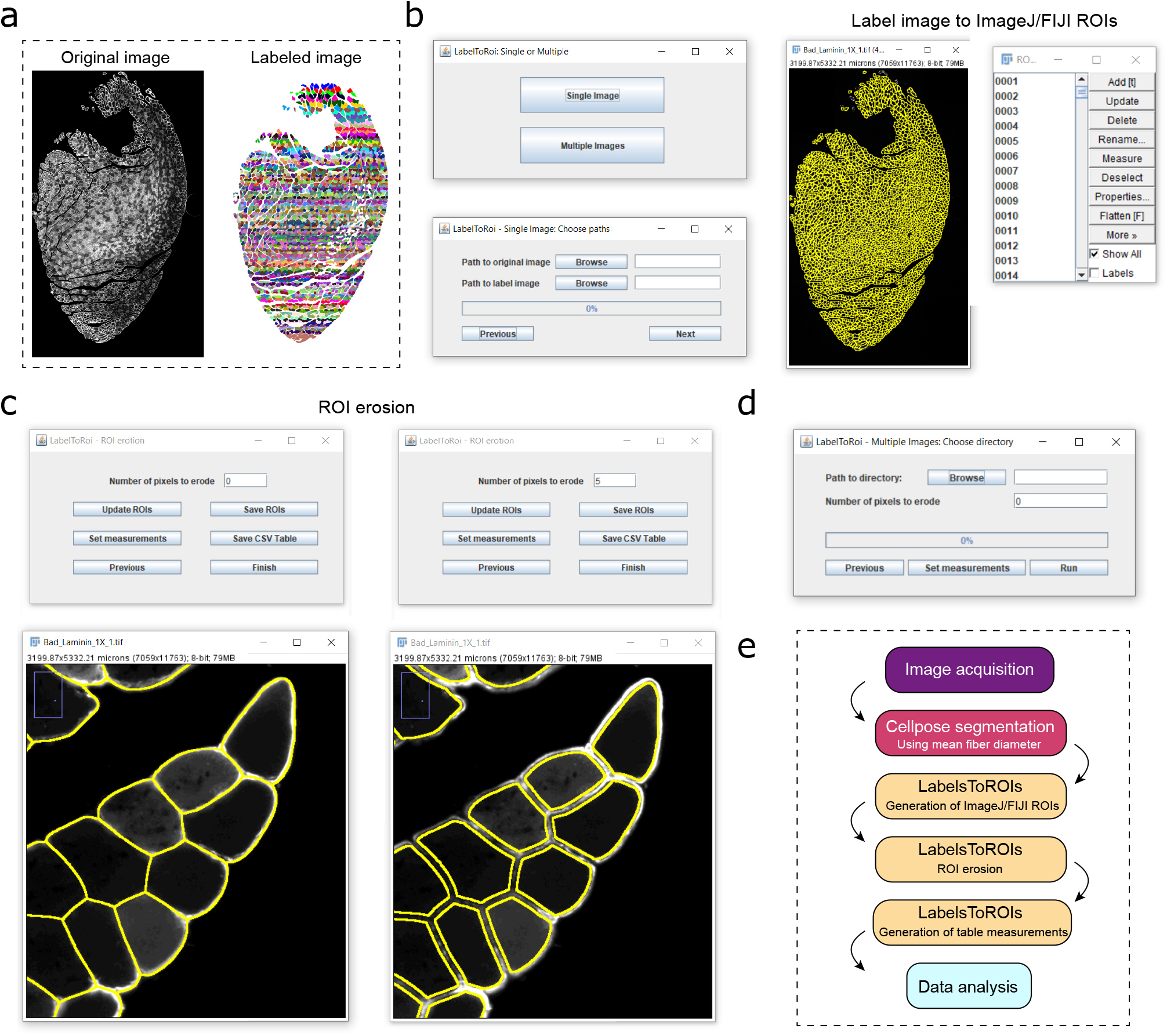
Development of LabelsToROIs plugin to analyze label images in FIJI/ImageJ. As indicated in (a), the plugin takes as input an original image and a labeled image generated using Cellpose or another software. (b) The plugin allows to automatically extract the segmentations in the label images and generate the corresponding FIJI/ImageJ regions of interest (ROIs). (c) Once generated, the ROIs can be eroded by a fixed number of pixels until the segmentations correctly delineate the objects of interest. The ROI files can be saved and different measurements can be selected to generate table measurements. (d) The LabelsToROIs plugin also allows for simultaneous processing of multiple images, enabling the generation of ROIs, ROI erosion and table measurements in one step, while automatically saving all the files for future inspection. (e) A diagram of the proposed pipeline for the automatic analysis of skeletal muscle biopsies.

### Cellpose combined with LabelsToRois allows for rapid and rigorous quantifications of myofibers in different injury and disease paradigms

After establishing that Cellpose can efficiently segment close to 100% of myofibers present in a cross-sections and the development of our ImageJ plugin to rapidly quantify myofibers in batch, we evaluated the use of Cellpose and LabelsToRois in different models of skeletal muscle regeneration and disease. First, we focused on a mouse model of Duchene muscular dystrophy (mdx mice), where intramuscular fat and fibrotic scar tissue gradually replace functional muscle^6^,48,49. We previously discovered that genetic removal of FAP cilia in mdx mice prevented fat formation^6^. Interestingly, we also found that loss of FAP cilia prevented the decline in myofiber size normally seen in mdx mice^6^. To test if Cellpose could generate comparable results, we used our original raw images from this study for Cellpose to segment (Fig. 7a). Validating our previous results, quantifications of the cross-sectional area of myofibers segmented by Cellpose and quantified by LabelToRois demonstrate that loss of FAP cilia in mdx mice (mdx FAPno cilia) efficiently prevented the decline in myofiber CSA (Fig. 7b) with a shift from fewer smaller to larger fibers (Fig. 7c) compared to mdx control mice. Besides reducing user bias, we also found that Cellpose was able to segment on average 400 more fibers per image compared to using manual thresholding. As a result, we saw an improvement in our statistical analysis by a factor of 10. Importantly, the use of LabelsToROIs plugin allowed to easily process all the labeled images in a single step and to generate a unique measurement table with all the quantifications in a tidy data format, greatly enhancing the subsequent analysis.

**Figure 7.**
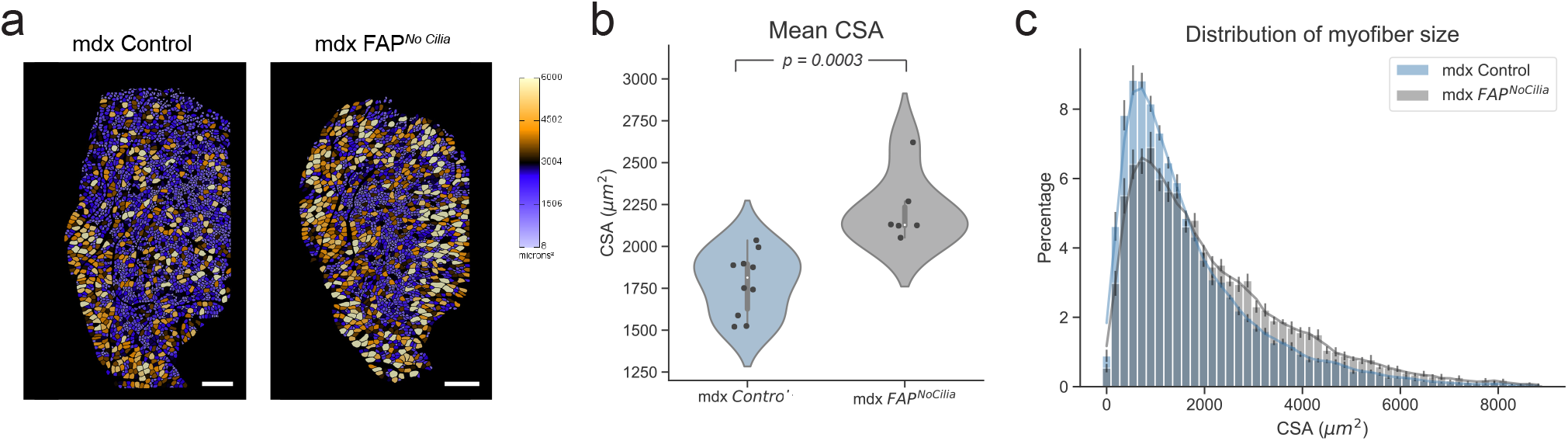
Cellpose and LabelsToROIs allow to faithfully quantify myofibers in a Duchene muscular dystrophy model. (a) Re-quantification of data from Kopinke et al., 2017. TA images from 10-to 12-month-old mdx control and mdx FAPNoCilia mice stained for LAMININ were re-analyzed using Cellpose. LabelsToROIs was used to erode the segmentations according to visual inspection (5 pixels). Myofiber segmentations were color-coded based on their CSA using the FIJI/ImageJ ROI color coder. Scale bar, 500 µm. (b) Mean CSA of mdx control (n=10) and mdx FAPNoCilia mice (n=6). Statistical differences were assessed by a Student T test. (c) Distribution of myofiber CSA for mdx control and mdx FAPNoCilia mice. Results are presented as means ± standard error of the mean (SEM) for each bin and each genotype.

We next evaluated the performance of Cellpose and LabelsToRois on detecting a difference in myofiber regeneration after different injury settings. We compared cardiotoxin (CTX) injury, the “gold-standard” muscle injury model^11,27,43,50^, which is rapidly repaired and causes little fat, to a glycerol (GLY) injury in CD1 wild type mice, which also elicits a regenerative response but results in massive fat infiltration^5,6,11,28,42–44^,. Confirming the differences in fat formation between these two injuries, we detected few PERILPIN+ fat cells 21 days post CTX injury (Fig. 8a) with around 25 adipocytes per mm^2^ of injured area (Fig. 8b). In contrast, GLY induced five times as much adipocyte differentiation as CTX, which persisted for up to 3 months post-GLY injury (Fig. 8a,b). We then asked if there is a difference in the regenerative response of myofibers between CTX and GLY at 21 days post-injury. Again, using Cellpose and our LabelsToRois plugin to automatically segment and quantify myofibers stained for LAMININ, we found that myofibers recovered more efficiently after a CTX injury compared to GLY (Fig. 8c,d). In fact, we noticed that a large majority of myofibers remained small compared to myofibers after a CTX injury (Fig. 8e). Thus, GLY causes massive fat infiltration and displays diminished myofiber regeneration capability compared to CTX. Our data further highlight the advantages of the GLY injury model as a valuable tool to mimic and study chronic fat infiltration, a hallmark of muscle disease and age, by simple intramuscular injections into any genetic background.

**Figure 8.**
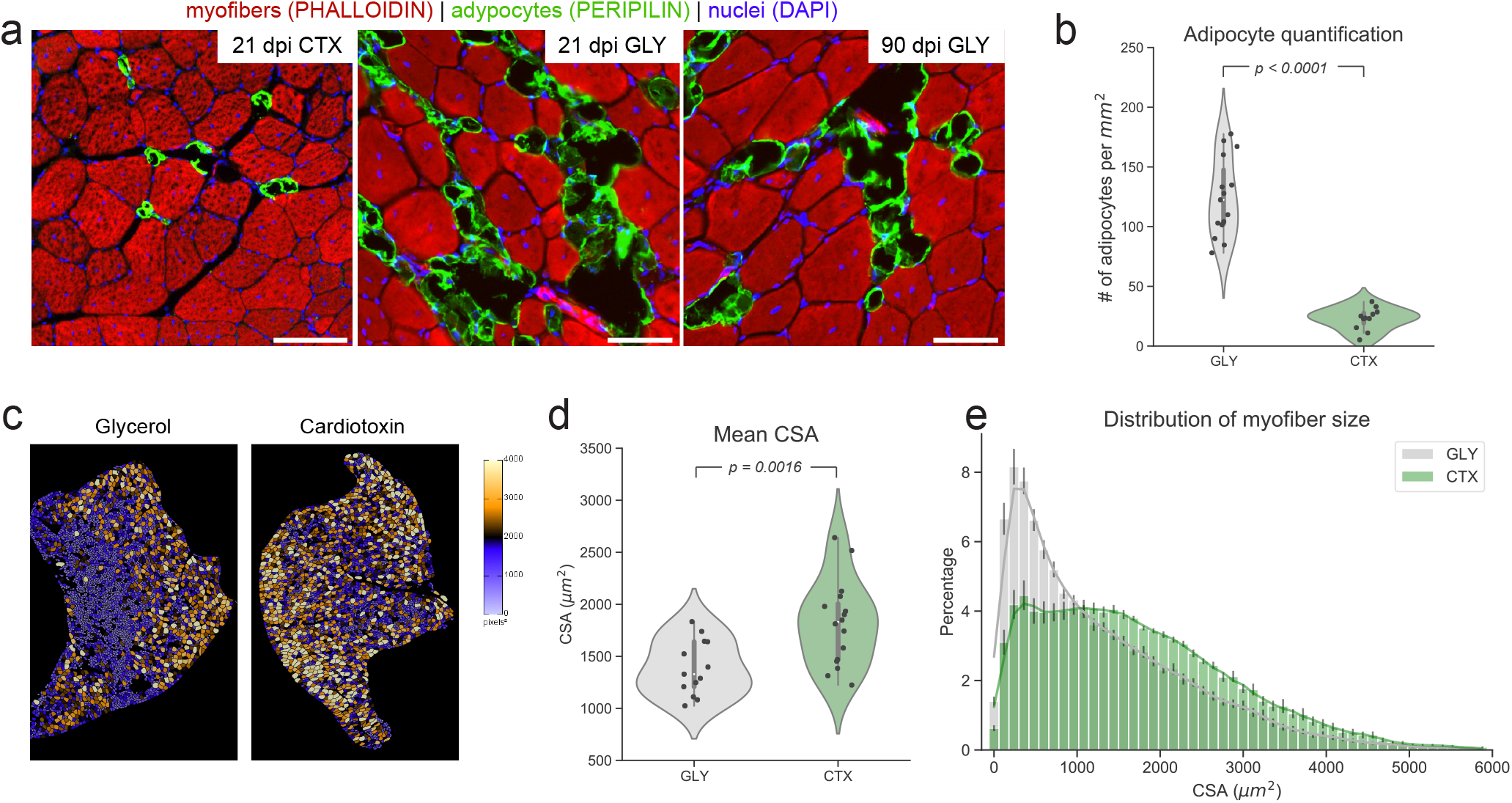
Cellpose and LabelsToROIs enable the quantification of regenerative recovery in different injury settings. (a)Few adipocytes, marked by PERILPIN (green), have formed 21 days post muscle injury caused by cardiotoxin (CTX). In contrast, glycerol-induced injury (GLY) causes massive fat infiltration that persists up to 3 months. Myofibers are labeled with Phalloidin (red) and nuclei with DAPI (blue). (b) Number of adipocytes per mm2 of injured area 21 days post GLY (n=15) or CTX (n=11) injury. Statistical differences were assessed by a Student T test. (c) Cellpose and LabelsToROIs were used to segment and process TA images stained for LAMININ corresponding to GLY and CTX treated mice. Myofiber segmentations were color-coded based on their CSA using the FIJI/ImageJ ROI color coder. (d) Quantification of the mean CSA of myofibers 21 days post GLY (n=12) or CTX (n=18) injury. Statistical differences were assessed by a Student T test. (c) Distribution of myofiber size for each injury. Results are presented as means ± standard error of the mean (SEM) for each bin and each treatment.

Finally, we evaluated the use of Cellpose and LabelsToROIs in determining myofiber types. Murine myofibers can be separated in slow twitch (type I) and fast twitch (type IIa, IIb & IIx) fibers and each muscle group is composed of a different combination of type I and type II fibers^51^. This composition can be altered through fiber type switching, caused by aging, exercise or disease^52^–59. Therefore, myofiber typing is an important assay to determine muscle health and regeneration. To determine if our segmentation pipeline can also be used for efficient myofiber typing, we stained muscle sections for type I and type IIa fibers as well as LAMININ to mark each myofiber. After segmenting the LAMININ channel with Cellpose, LabelsToROIs was used for label erosion and fluorescence intensity quantification in the different channels. This analysis allowed us to automatically quantify the total number of fibers present as well as their corresponding CSAs (Fig. S2). Alternatively, Cellpose can be used to individually segment the images corresponding to each stain, allowing a rapid evaluation of the proportion of fiber types and their CSA. Thus, co-staining muscle section with LAMININ and fiber type specific antibodies allows for rapid determination of the number, type and CSA of all myofibers present in a given muscle section.

## Conclusion

Myofiber CSA is a crucial measurement to assess muscle regeneration as well as degeneration. While several CSA plugins have been developed to quantify myofiber CSA in an automated and unbiased fashion, including the ability to quantify a variety of cell types, they all require “perfect staining”. The biggest advantage of Cellpose is that it can be used to segment suboptimal stainings obtained from fixed tissue with remarkable high fidelity. As reported here, we found that Cellpose was equally able to segment myofibers marked by LAMININ, Phalloidin and WGA. Thus, Cellpose is able to segment myofibers labeled by a variety of markers. Importantly, we found that although Cellpose accurately identified the majority of the myofibers, there was an area bias associated to the type of staining used, which significantly affected myofiber CSA. The development of the ImageJ plugin LabelsToROIs allowed us to solve this issue, enabling the identifation of myofibers with an accuracy similar to human experts. Thus, the combined use of the novel deep learning algorithm Cellpose and our LabelsToRois plugin allow for unbiased and rigorous analysis of thousands of myofibers within minutes even by users not well-versed in computer programming, regardless of staining or fixation condition. Importantly, the use of this pipeline is not limited to skeletal muscle analysis and can be extended to different experimental setups. Thus, we believe that our pipeline will reduce bias, enhance reproducibility, and allow widespread use of this segmentation tool.

## Methods

### Animal studies

Ift88tm1Bky and (Tg(Pdgfr-cre/ERT)467Dbe) alleles have been described previously^60,61^. Mdx mice were purchased from the Jackson Laboratory (Stock No: 001801). Littermates lacking either the Cre or the null (in the case of Ift88) served as controls. Tamoxifen (Sigma T-5648) was dissolved in corn oil and administered by oral gavage (200-250 mg/kg) on two consecutive days to all mice at 4-5 weeks of age. Muscle tissue was then harvested at 1 year of age. For Cardiotoxin and Glycerol injuries, male and female wild type mice on a mixed CD1 background at 10-12 weeks of age were used. To evaluate muscle regeneration, mice were anesthetized with isofluorane and the Tibialis Anterior (TA) was injected with 25-50µL of either 10µM Cardiotoxin (Naja Pallida, Sigma) or 50% Glycerol. The TA was allowed to regenerate for 21 days before TAs were harvested. All animal protocols were approved by the Institutional Animal Care and Use Committee (IACUC) of the University of Florida. Histology, Immunohistochemistry and Image Analysis Tissue fixation, processing and immunostaining were performed as described previously^6,40^. In brief, TAs were fixed or snap frozen in order to compare antibody stainings and subsequent segmentation analysis. For tissue fixation, the TAs were submerged in paraformaldehyde (PFA) 4% for 3hrs at 4°C, washed three times for 5min in cold PBS, kept overnight in sucrose 30% before embedded in OCT, and frozen in liquid nitrogen-cooled isopentane. To snap freeze, TAs were immediately embedded in OCT after harvesting, and frozen in liquid nitrogen-cooled isopentane. TAs were cryosectioned at midbelly at 10µm, multiple sections were collected and stored at-80°C until staining. For immunofluorescent staining, slides were thawed, washed in PBST (0.1% Tween-20 in PBS) three times for 5min and incubated overnight at 4°C in blocking solution (5% donkey serum in PBS with 0.3%Triton X-100). Primary antibodies were incubated overnight at 4°C, followed by three 5min washes in PBST. Slides were then incubated with secondary antibodies along with directly conjugated probes for 45min, washed three times for 5min each and mounted with Fluoromount-G (ThermoFisher). Muscle fibers were visualized with the directly conjugated probe Phalloidin-Alexa-568 (1:250, Molecular Probes A12380). Fibers were additionally outlined by marking the extracellular matrix with both the directly conjugated wheat germ agglutinin (WGA)-Alexa 647 (Molecular Pobes, W32466), and the primary antibody Rabbit anti-LAMININ (1:1000, Sigma L9393) and the corresponding donkey antirabbit Alexa Fluor 488 (1:1000, A21206). For fiber type analysis, snap frozen sections were used, mouse-on-mouse blocking was performed with AffiniPure Fab Fragment Donkey Anti-Mouse (1:50, Jackson ImmunoResearch, 715-007-003) during the blocking step. The following primary antibodies were used: Myosin heavy chain Type 1 (1:50, supernatant from DSHB, BA-D5), Myosin heavy chain Type IIA (1:50, supernatant from DSHB, SC-71), along with LAMININ to delineate all fibers. Secondaries were goat anti-mouse IgG Fc subclass 2b specific Alexa Fluor 488 (1:800, Jackson ImmunoResearch, 115-545-207), donkey antirabbit Alexa Fluor 568 (1:1000, Invitrogen, A10042), and goat anti-mouse Alexa Fluor 647 IgG Fc subclass 1 specific (1:800, Jackson ImmunoResearch 115-605-205), respectively. To visualize primary cilia, FAPs and adipocytes, the following primary antibodies were used: anti-ARL13B (1:1000, Proteintech 17711-1-AP), goat anti-PDGFR (1:250, RD Systems AF1062) and abbit anti-PERILIPIN (1:1000, Cell Signaling 9349). Species-correspodning Alexa Fluor-conjugated secondary antibodies from Life Technologies (1:1000) were used and DAPI (Sigma) to visualize nuclei. Images of the most representative section were acquired using a Leica TCS SPE confocal equipped with a DFC7000 camera. The LAS Navigator software was used to generate a merged image of the whole TA cross-section. All images were processed identically with Adobe Photoshop (CS5) or ImageJ.

### Development of LabelsToRois plugin and tutorials

LabelsToRois plugin was developed in Jython. The plugin, together with a detailed user tutorial which includes Cellpose usage, can be accessed from the following link: https://labelstorois.github.io/

### Image analysis

Single channel fluorescence images were segmented with Cellpose using either the GUI or with a modified version from the Python script provided by the authors. Labeled images resulting from Cellpose segmentation were fed to the ImageJ plugin LabelsToRois together with the original image. The resulting ROIs were eroded with a fixed number of pixels according to visual inspection. Area measurements were generated and analyzed with custom Python scripts.

Dice coefficient was calculated for each pair of labeled images using the SimpleITK library in Python^62^. For that, labels IDs were previously matched using custom scripts. To compute the Dice coefficient for the eroded labels, ImageJ ROIs were eroded using the LabelsToROIs plugin and then labeled images we generated in ImageJ using the ROI map function from the LOCI update site. Connected components labeling and Set Label Map functions from the MorpholibJ plugin were also used for working with labeled images^63^.

## Acknowledgements

The authors’ work presented in this work was supported a departmental start-up funds to DK. AW is supported by CONICET. AMN is supported by a National Institute of Child Health and Human Development Grant (T32-HD-043730). We thank Ambili Appu and Connor Johnson for technical help with data aquisition. We are also thankful for critical input to Santiago Miriuka, members of the Miriuka Lab, and members of the Myology Institute at the University of Florida on the manuscript. Figure illustrations were drawn by AW.

## Author contributions

Study design (AW, AMN DK), Experimental procedures, data acquisition and analysis (AW, AMN MEC). All authors were involved in data interpretation, data presentation, and manuscript writing.

## Supplementary Figures

**Figure S1.**
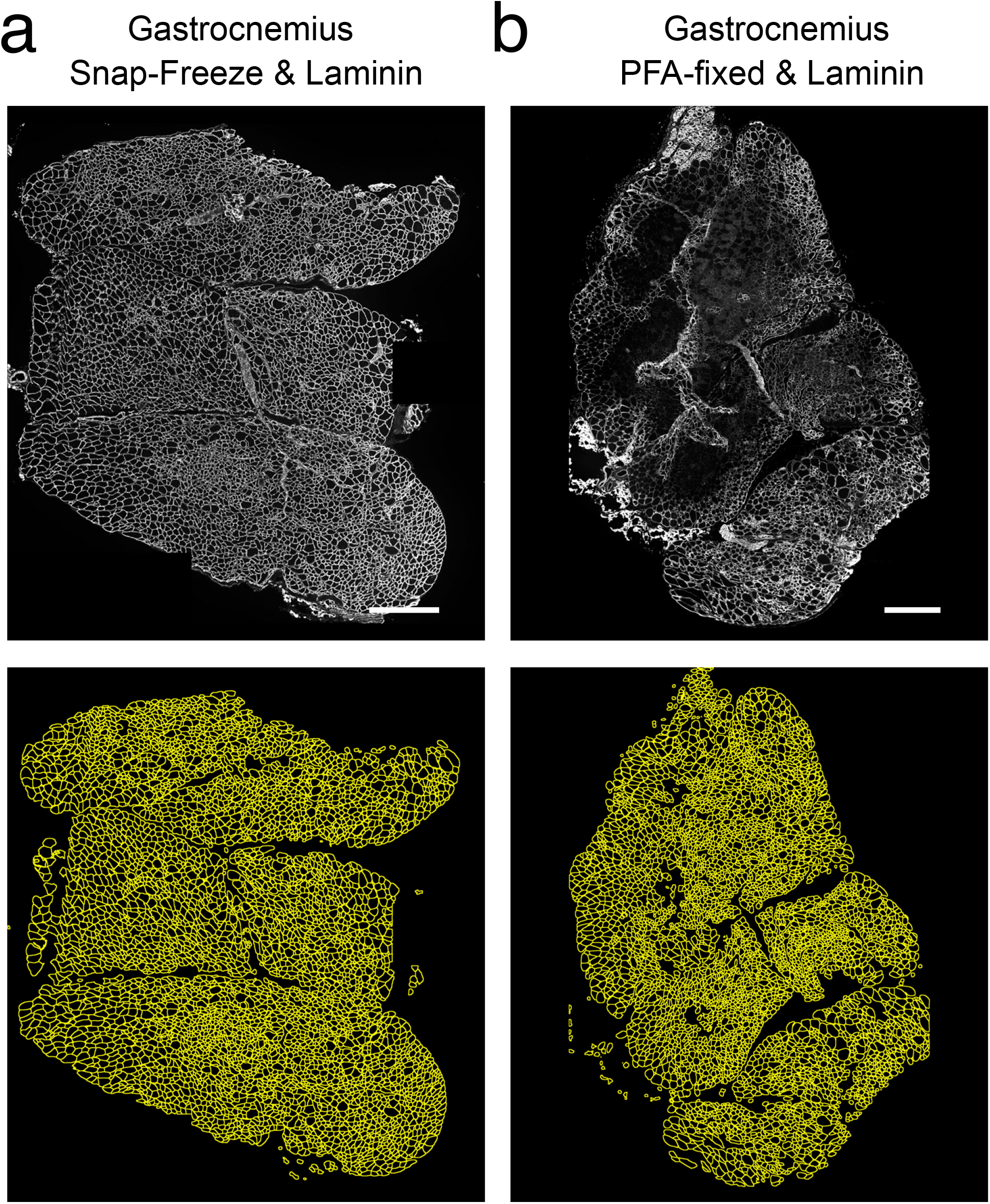
Cellpose segmentation of gastrocnemius muscle. Cross section of (a) snap-frozen or (b) PFA-fixed gastrocnemius muscle stained for LAMININ. Top, original image. Bottom, Cellpose segmentation results. Scale bar, 500 µm.

**Figure S2.**
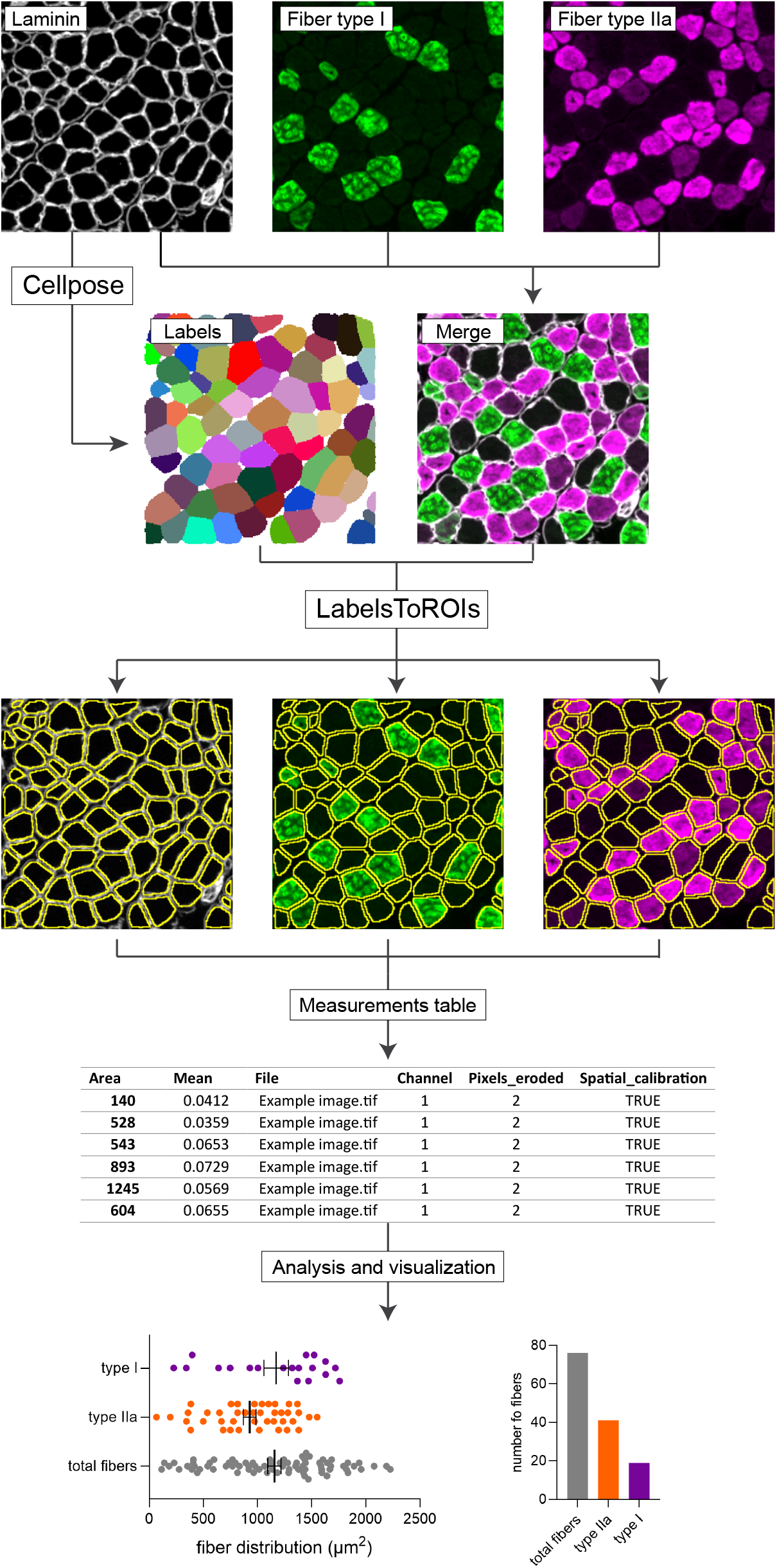
Segmentation pipeline enables simple and automated myofiber type analysis. Snap frozen gastrocnemius muscle section were stained with LAMININ (gray scale) to delineate all fibers, BA-D5 to mark Type 1 fibers (green) and SC-71 for Type IIA fibers (purple). The LAMININ image was fed to Cellpose for myofiber segmentation. The resulting labeled image, together with the merged image containing all the channels were fed to the LabelsToROIs plugin for ROI erosion and fluorescence intensity quantification. Analysis of the resulting table in Excel allowed the calculation of myofiber numbers and size distribution.

## References

1. Lepper, C., Partridge, T. A. & Fan, C. M. An absolute requirement for pax7-positive satellite cells in acute injury-induced skeletal muscle regeneration. Development 138, 3639–3646, DOI: 10.1242/dev.067595 (2011).

2. Murphy, M. M., Lawson, J. A., Mathew, S. J., Hutcheson, D. A. & Kardon, G. Satellite cells, connective tissue fibroblasts and their interactions are crucial for muscle regeneration. Development DOI: 10.1242/dev.064162 (2011).

3. Relaix, F. & Zammit, P. S. Satellite cells are essential for skeletal muscle regeneration: the cell on the edge returns centre stage. Development DOI: 10.1242/dev.069088 (2012).

4. Sambasivan, R. et al. Pax7-expressing satellite cells are indispensable for adult skeletal muscle regeneration. Development 138, 3647–3656, DOI: 10.1242/dev.067587 (2011).

5. Joe, A. W. et al. Muscle injury activates resident fibro/adipogenic progenitors that facilitate myogenesis. Nat. Cell Biol. 12, DOI: 10.1038/ncb2015 (2010).

6. Kopinke, D., Roberson, E. C. & Reiter, J. F. Ciliary Hedgehog Signaling Restricts Injury-Induced Adipogenesis. Cell 170, 340–351.e12, DOI: 10.1016/j.cell.2017.06.035 (2017).

7. Lukjanenko, L. et al. Aging Disrupts Muscle Stem Cell Function by Impairing Matricellular WISP1 Secretion from Fibro-Adipogenic Progenitors. Cell Stem Cell 24, 433–446.e7, DOI: 10.1016/j.stem.2018.12.014 (2019).

8. Murphy, M. & Kardon, G. Origin of vertebrate limb muscle: The role of progenitor and myoblast populations, vol. 96 (2011).

9. Santini, M. P. et al. Tissue-Resident PDGFRα+ Progenitor Cells Contribute to Fibrosis versus Healing in a Context- and Spatiotemporally Dependent Manner. Cell Reports 30, 555–570.e7, DOI: 10.1016/j.celrep.2019.12.045 (2020).

10. Uezumi, A., Fukada, S. I., Yamamoto, N., Takeda, S. & Tsuchida, K. Mesenchymal progenitors distinct from satellite cells contribute to ectopic fat cell formation in skeletal muscle. Nat. Cell Biol. 12, DOI: 10.1038/ncb2014 (2010).

11. Uezumi, A. et al. Fibrosis and adipogenesis originate from a common mesenchymal progenitor in skeletal muscle. J. Cell Sci. 124, DOI: 10.1242/jcs.086629 (2011).

12. Wosczyna, M. N. et al. Mesenchymal Stromal Cells Are Required for Regeneration and Homeostatic Maintenance of Article Mesenchymal Stromal Cells Are Required for Regeneration and Homeostatic Maintenance of Skeletal Muscle. CellReports 27, 2029–2035.e5, DOI: 10.1016/j.celrep.2019.04.074 (2019).

13. Hogarth, M. W. et al. Fibroadipogenic progenitors are responsible for muscle loss in limb girdle muscular dystrophy 2B. Nat. Commun. 10, 1–13, DOI: 10.1038/s41467-019-10438-z (2019).

14. Liu, W., Liu, Y., Lai, X. & Kuang, S. Intramuscular adipose is derived from a non-Pax3 lineage and required for efficient regeneration of skeletal muscles. Dev. Biol. DOI: 10.1016/j.ydbio.2011.10.011 (2012).

15. Scott, R. W., Arostegui, M., Schweitzer, R., Rossi, F. M. & Underhill, T. M. Hic1 Defines Quiescent Mesenchymal Progenitor Subpopulations with Distinct Functions and Fates in Skeletal Muscle Regeneration. Cell Stem Cell 25, 797–813.e9, DOI: 10.1016/j.stem.2019.11.004 (2019).

16. Stumm, J. et al. Odd skipped-related 1 (Osr1) identifies muscle-interstitial fibro-adipogenic progenitors (FAPs) activated by acute injury. Stem Cell Res. 32, 8–16, DOI: 10.1016/j.scr.2018.08.010 (2018).

17. Burakiewicz, J. et al. Quantifying fat replacement of muscle by quantitative MRI in muscular dystrophy. J. Neurol. 264, 2053–2067, DOI: 10.1007/s00415-017-8547-3 (2017).

18. Goodpaster, B. H. et al. Obesity, regional body fat distribution, and the metabolic syndrome in older men and women. Arch. Intern. Medicine 165, 777–783, DOI: 10.1001/archinte.165.7.777 (2005).

19. Goodpaster, BH; Krishnaswami, S. et al. Tissue Distribution and Both Type 2 Diabetes and Impaired Glucose. Diabetes care 26, 372–379 (2003).

20. Goodpaster, B. H., Thaete, F. L. & Kelley, D. E. Thigh adipose tissue distribution is associated with insulin resistance in obesity and in type 2 diabetes mellitus. Am. J. Clin. Nutr. 71, 885–892, DOI: 10.1093/ajcn/71.4.885 (2000).

21. Goodpaster, B. H. et al. The loss of skeletal muscle strength, mass, and quality in older adults: The Health, Aging and Body Composition Study. Journals Gerontol. - Ser. A Biol. Sci. Med. Sci. 61, 1059–1064, DOI: 10.1093/gerona/61.10.1059 (2006).

22. Goodpaster, B. H., Theriault, R., Watkins, S. C. & Kelley, D. E. Intramuscular lipid content is increased in obesity and decreased by weight loss. Metab. Clin. Exp. 49, 467–472, DOI: 10.1016/S0026-0495(00)80010-4 (2000).

23. Milad, N. et al. Increased plasma lipid levels exacerbate muscle pathology in the mdx mouse model of Duchenne muscular dystrophy. Skeletal Muscle 7, 1–14, DOI: 10.1186/s13395-017-0135-9 (2017).

24. Murphy, W. A., Totty, W. G. & Carroll, J. E. MRI of normal and pathologic skeletal muscle. Am. J. Roentgenol. 146, 565–574, DOI: 10.2214/ajr.146.3.565 (1986).

25. Willcocks, R. J. et al. Multicenter prospective longitudinal study of magnetic resonance biomarkers in a large duchenne muscular dystrophy cohort. Annals Neurol. 79, 535–547, DOI: 10.1002/ana.24599 (2016).

26. Wokke, B. H. et al. Quantitative MRI and strength measurements in the assessment of muscle quality in Duchenne muscular dystrophy. Neuromuscul. Disord. 24, 409–416, DOI: 10.1016/j.nmd.2014.01.015 (2014).

27. Hardy, D. et al. Comparative Study of Injury Models for Studying Muscle Regeneration in Mice. PLOS ONE 11, e0147198, DOI: 10.1371/journal.pone.0147198 (2016).

28. Mahdy, M. A., Lei, H. Y., Wakamatsu, J. I., Hosaka, Y. Z. & Nishimura, T. Comparative study of muscle regeneration following cardiotoxin and glycerol injury. Annals Anat. DOI: 10.1016/j.aanat.2015.07.002 (2015).

29. Miller, M. S., Bedrin, N. G., Ades, P. A., Palmer, B. M. & Toth, M. J. Molecular determinants of force production in human skeletal muscle fibers: Effects of myosin isoform expression and cross-sectional area. Am. J. Physiol. - Cell Physiol. 308, C473–C484, DOI: 10.1152/ajpcell.00158.2014 (2015).

30. Virgilio, K. M., Martin, K. S., Peirce, S. M. & Blemker, S. S. Agent-based model illustrates the role of the microenvironment in regeneration in healthy and mdx skeletal muscle. J. Appl. Physiol. 125, 1424–1439, DOI: 10.1152/japplphysiol.00379.2018 (2018).

31. Briguet, A., Courdier-Fruh, I., Foster, M., Meier, T. & Magyar, J. P. Histological parameters for the quantitative assessment of muscular dystrophy in the mdx-mouse. Neuromuscul. Disord. 14, 675–682, DOI: 10.1016/j.nmd.2004.06.008 (2004).

32. Duddy, W. et al. Muscular dystrophy in the mdx mouse is a severe myopathy compounded by hypotrophy, hypertrophy and hyperplasia. Skeletal Muscle 5, 1–18, DOI: 10.1186/s13395-015-0041-y (2015).

33. Flück, M. Functional, structural and molecular plasticity of mammalian skeletal muscle in response to exercise stimuli. J. Exp. Biol. 209, 2239–2248, DOI: 10.1242/jeb.02149 (2006).

34. McDonald, A. A., Hebert, S. L., Kunz, M. D., Ralles, S. J. & McLoon, L. K. Disease course in mdx:Utrophin+/ mice: comparison of three mouse models of duchenne muscular dystrophy. Physiol. Reports 3, 1–22, DOI: 10.14814/phy2.12391 (2015).

35. Desgeorges, T. et al. Open-CSAM, a new tool for semi-automated analysis of myofiber cross-sectional area in regenerating adult skeletal muscle. Skeletal Muscle 9, DOI: 10.1186/s13395-018-0186-6 (2019).

36. Encarnacion-Rivera, L., Foltz, S., Hartzell, H. C. & Choo, H. Myosoft: An automated muscle histology analysis tool using machine learning algorithm utilizing FIJI/ImageJ software. PLoS ONE 15, 1–22, DOI: 10.1371/journal.pone.0229041 (2020).

37. Kastenschmidt, J. M. et al. QuantiMus: A Machine Learning-Based Approach for High Precision Analysis of Skeletal Muscle Morphology. Front. Physiol. 10, DOI: 10.3389/fphys.2019.01416 (2019).

38. Reyes-Fernandez, P. C., Periou, B., Decrouy, X., Relaix, F. & Authier, F. J. Automated image-analysis method for the quantification of fiber morphometry and fiber type population in human skeletal muscle. Skeletal Muscle 9, 1–15, DOI: 10.1186/s13395-019-0200-7 (2019).

39. Smith, L. R. & Barton, E. R. SMASH - semi-automatic muscle analysis using segmentation of histology: A MAT-LAB application. Skeletal Muscle 4, DOI: 10.1186/2044-5040-4-21 (2014).

40. Hilgendorf, K. I. et al. Omega-3 Fatty Acids Activate Ciliary FFAR4 to Control Adipogenesis Article Omega-3 Fatty Acids Activate Ciliary FFAR4 to Control Adipogenesis. Cell 179, 1289–1305.e21, DOI: 10.1016/j.cell.2019.11.005 (2019).

41. Stringer, C., Wang, T., Michaelos, M. & Pachitariu, M. Cellpose: a generalist algorithm for cellular segmentation. Nat. Methods 18, 100–106, DOI: 10.1038/s41592-020-01018-x (2021).

42. Kawai, H. et al. Experimental glycerol myopathy: a histological study. Acta Neuropathol. 80, 192–197, DOI: 10.1007/BF00308923 (1990).

43. Lukjanenko, L., Brachat, S., Pierrel, E., Lach-Trifilieff, E. & Feige, J. N. Genomic Profiling Reveals That Transient Adipogenic Activation Is a Hallmark of Mouse Models of Skeletal Muscle Regeneration. PLoS ONE 8, e71084, DOI: 10.1371/journal.pone.0071084 (2013).

44. Pisani, D. F., Bottema, C. D., Butori, C., Dani, C. & Dechesne, C. A. Mouse model of skeletal muscle adiposity: A glycerol treatment approach. Biochem. Biophys. Res. Commun. DOI: 10.1016/j.bbrc.2010.05.021 (2010).

45. Carass, A. et al. Evaluating White Matter Lesion Segmentations with Refined Sørensen-Dice Analysis. Sci. Reports 10, DOI: 10.1038/s41598-020-64803-w (2020).

46. Dice, L. R. Measures of the Amount of Ecologic Association Between Species Author (s): Lee R. Dice Published by: Ecological Society of America Stable URL: http://www.jstor.org/stable/1932409. Ecology 26, 297–302 (1945).

47. Zou, K. H. et al. Statistical Validation of Image Segmentation Quality Based on a Spatial Overlap Index. Acad. Radiol. 11, 178–189, DOI: 10.1016/S1076-6332(03)00671-8 (2004).

48. Bulfield, G., Siller, W. G., Wight, P. A. & Moore, K. J. X chromosome-linked muscular dystrophy (mdx) in the mouse. Proc. Natl. Acad. Sci. United States Am. 81, 1189– 1192, DOI: 10.1073/pnas.81.4.1189 (1984).

49. Hoffman, E. P., Brown, R. H. & Kunkel, L. M. Dystrophin: The protein product of the duchenne muscular dystrophy locus. Cell 51, 919–928, DOI: 10.1016/0092-8674(87)90579-4 (1987).

50. Ownby, C. L., Fletcher, J. E. & Colberg, T. R. Cardiotoxin 1 from cobra (Naja naja atra) venom causes necrosis of skeletal muscle in vivo. Toxicon 31, 697–709, DOI: 10.1016/0041-0101(93)90376-T (1993).

51. Schiaffino, S. & Reggiani, C. Fiber types in Mammalian skeletal muscles. Physiol. Rev. 91, 1447–1531, DOI: 10.1152/physrev.00031.2010 (2011).

52. Evans, W. J. & Lexell, J. Human Aging, Muscle Mass, and Fiber Type Composition. The Journals Gerontol. Ser. A 50A, 11–16, DOI: 10.1093/gerona/50A.Special_Issue.11 (1995).

53. Handschin, C. et al. Skeletal muscle fiber-type switching, exercise intolerance, and myopathy in PGC-1α muscle-specific knock-out animals. J. Biol. Chem. 282, 30014– 30021, DOI: 10.1074/jbc.M704817200 (2007).

54. Rafael, J. A. et al. Dystrophin and utrophin influence fiber type composition and post-synaptic membrane structure. Hum. Mol. Genet. 9, 1357–1367, DOI: 10.1093/hmg/9.9.1357 (2000).

55. Reyes, N. L. et al. Fnip1 regulates skeletal muscle fiber type specification, fatigue resistance, and susceptibility to muscular dystrophy. Proc. Natl. Acad. Sci. United States Am. 112, 424–429, DOI: 10.1073/pnas.1413021112 (2015).

56. Röckl, K. S. et al. Skeletal muscle adaptation to exercise training: AMP-activated protein kinase mediates muscle fiber type shift. Diabetes 56, 2062–2069, DOI: 10.2337/db07-0255 (2007).

57. Selsby, J. T., Morine, K. J., Pendrak, K., Barton, E. R. & Sweeney, H. L. Rescue of dystrophic skeletal muscle by PGC-1α involves a fast to slow fiber type shift in the mdx mouse. PLoS ONE 7, 1–10, DOI: 10.1371/journal.pone.0030063 (2012).

58. Talbot, J. & Maves, L. Skeletal muscle fiber type: using insights from muscle developmental biology to dissect targets for susceptibility and resistance to muscle disease. Wiley Interdiscip. Rev. Dev. Biol. 5, 518–534, DOI: 10.1002/wdev.230 (2016).

59. Thompson, L. D. V. Skeletal muscle adaptations with age, inactivity, and therapeutic exercise. J. Orthop. Sports Phys. Ther. 32, 44–57, DOI: 10.2519/jospt.2002.32.2.44 (2002).

60. Haycraft, C. J. et al. Intraflagellar transport is essential for endochondral bone formation. Development 134, 307– 316, DOI: 10.1242/dev.02732 (2007).

61. Kang, S. H., Fukaya, M., Yang, J. K., Rothstein, J. D. & Bergles, D. E. NG2+ CNS glial progenitors remain committed to the oligodendrocyte lineage in postnatal life and following neurodegeneration. Neuron 68, 668–681, DOI: 10.1016/j.neuron.2010.09.009 (2010).

62. Lowekamp, B. C., Chen, D. T., Ibáñez, L. & Blezek, D. The design of simpleITK. Front. Neuroinformatics 7, DOI: 10.3389/fninf.2013.00045 (2013).

63. Legland, D., Arganda-Carreras, I. & Andrey, P. MorphoLibJ: Integrated library and plugins for mathematical morphology with ImageJ. Bioinformatics 32, DOI: 10.1093/bioinformatics/btw413 (2016).

